# Pumping the brakes: rostromedial tegmental inhibition of compulsive cocaine seeking

**DOI:** 10.1101/2023.10.04.560908

**Authors:** Peter J. Vento, Jacob R. Watson, Dominika Pullmann, Samantha L. Black, Jensen S. Tomberlin, Thomas C. Jhou

## Abstract

Addiction is marked by aberrant decision-making and an inability to suppress inappropriate and often dangerous behaviors. We previously demonstrated that inactivation of the rostromedial tegmental nucleus (RMTg) in rats causes persistent food seeking despite impending aversive footshock, an effect strikingly similar to the punishment resistance observed in people with a history of protracted drug use [1]. Here, we extend these studies to demonstrate chemogenetic silencing of RMTg axonal projections to the ventral tegmental area (VTA) (RMTg→VTA pathway) causes rats to endure significantly more footshock to receive cocaine infusions. To further test whether activation of this circuit is sufficient to suppress reward seeking in the absence of an overtly aversive stimulus, we used temporally specific optogenetic stimulation of the RMTg→VTA pathway as a “punisher” in place of footshock following lever pressing for either food or cocaine reward. While optical stimulation of the RMTg→VTA pathway robustly suppressed lever pressing for food, we found that stimulation of this circuit had only modest effects on suppressing responding for cocaine infusions. Even though optical RMTg→VTA stimulation was not particularly effective at reducing *ongoing* cocaine use, this experience nevertheless had long-lasting consequences, as reinstatement of drug seeking in response to cocaine-associated cues was profoundly suppressed when tested nearly two weeks later. These results suggest the RMTg may serve as a useful target for producing enduring reductions in drug craving, particularly during periods of abstinence from drug use.

## Introduction

Drug addiction is a chronic relapsing disorder that often leads to dangerous and exceedingly risky patterns of behavior that persist despite high monetary, physiological, and interpersonal costs. Still, more than 85% of people battling substance abuse relapse within their first year of treatment [2]. Long-term cocaine users, for example, exhibit impairments in inhibitory control and executive function [3], display perseverative behavior [4], and overvalue immediate rewards over larger delayed alternatives [1]. While evidence suggests important roles for mesolimbic dopamine (DA) [3, 5] and fronto-striatal networks [6, 7] in mediating pro-drug-seeking behaviors, investigation into aversive learning circuits and mechanisms inducing drug avoidance may hold the key to improving inhibitory control, reducing the risk of relapse and suppressing aberrant decision-making to improve long-term outcomes in substance use disorder.

The rostromedial tegmental nucleus (RMTg) provides dense GABAergic projections to midbrain DA neurons, and recent findings suggest an important role for the RMTg in behavioral inhibition and punishment learning [8–10]. The RMTg is activated by a wide range of aversive stimuli, including footshocks, shock-associated cues, reward omission, and drug withdrawal [11–14], and RMTg activation leads to place aversion [15] indicating a prominent RMTg role in both the encoding and expression of aversive responses. Our group has shown that optogenetic inhibition of RMTg projections to the ventral tegmental area (VTA) (RMTg→VTA pathway) robustly increases the intensity of footshock rats are willing to endure to receive food reward, reminiscent of resistance to punishment reported in humans and rodents after extended drug use [8]. Further, the RMTg has been shown to play a prominent role in aversive responses to cocaine [16], and pharmacological inhibition of the rostral tegmentum via infusion of the GABA agonists baclofen and muscimol caused increased lever pressing and impaired behavioral inhibition in a cued reinstatement test of cocaine seeking [17]. Still, the broader neurocircuitry through which the RMTg modulates drug seeking remains unclear, and the RMTg role in modulating drug seeking under punishment remains an open question.

To further explore the RMTg contribution to compulsive drug seeking, we modulated the RMTg→VTA pathway during two distinct addiction-related behaviors: persistent reward seeking under punishment and cue-induced reinstatement of cocaine seeking. Specifically, given previous data indicating that prolonged cocaine exposure induces resistance to the suppressive effect of footshock on lever pressing for cocaine [18, 19], we tested whether chemogenetic inhibition of the RMTg→VTA pathway results in a punishment-resistant phenotype when lever pressing for cocaine is immediately punished by brief footshock. Next, we tested whether timing- and pathway-specific optogenetic stimulation of the RMTg→VTA pathway, in the absence of an overtly aversive stimulus, is sufficient to suppress lever pressing for food or cocaine reward, or cue-induced reinstatement of cocaine seeking. Together, these data indicate the RMTg→VTA pathway is a critical mediator in suppressing drug use under punishment, and modulation of this circuit produces long-lasting reductions in cocaine seeking.

## Methods

### Animals

Adult male Sprague-Dawley rats weighing approximately 300g upon arrival from vendor (Charles River Laboratories, Raleigh, NC) were individually housed in standard shoebox cages in a temperature- and humidity-controlled vivarium with food and water provided *ad libitum*, unless stated otherwise. All procedures conformed to the Medical University of South Carolina and University of South Carolina *Institutional Animal Care and Use Committee* and the National Institutes of Health *Guide for the Care and Use of Laboratory Animals*.

### Drugs

Cocaine hydrochloride (HCl) and clozapine N-oxide (CNO) were provided by the National Institute of Drug Abuse (NIDA) Drug Supply Program (Research Triangle Park, NC, USA).

### Intravenous catheterization surgery and self-administration training

Rats were anesthetized using inhaled isoflurane and a chronic intravenous catheter was implanted in the jugular vein and passed subcutaneously to a guide cannula sutured through the animal’s back. Ketoprofen (5mg/kg, sc) was given during surgery and as needed post-surgery to reduce pain and swelling.

Catheters were flushed daily using sterile 0.9% saline and taurolidine-citrate solution (TCS, 0.05mL). At least 5 days after surgery, rats began training for intravenous cocaine self-administration. Training consisted of once-daily 2hr sessions during which time a single press on the active lever (FR1) initiated a light and tone cue and cocaine infusion (0.75 mg/kg/infusion, dissolved in 0.9% saline). Both the active (right) and inactive (left) levers remained extended for the duration of each session, and a house light was illuminated. A 20sec intertrial interval (ITI) was imposed during which time pressing on either lever yielded no consequences, thereby avoiding spurious infusions. Rats were considered to have successfully acquired self-administration after 10 consecutive days receiving ≥10 infusions. Extinction training consisted of 2hr sessions in which both the active and inactive levers were extended, and the house light illuminated; however, presses on the active lever did not result in drug delivery or presentation of the light/tone cues. Responses on the inactive lever did not result in any programmed consequences at any point in the experiment.

### Stereotaxic surgery

Under isoflurane anesthesia (5% for induction, 2% maintenance throughout surgery) rats were fixed in a stereotaxic frame, a small incision was made in the scalp, and small burr holes were drilled in the skull. Virus was injected into the RMTg (10° angle; from bregma AP: -7.4mm, ML: +1.9mm; DV: -7.4mm from dura) using Nanoject Auto-nanoliter Injectors (100nL/min; Drummond Scientific Company, Broomall, PA) with pulled glass pipettes. Injectors were left in place for at least 5 min post-infusion to permit diffusion of virus. Either guide cannulae (26 gauge, 11mm length; Plastics One, Roanoke, VA) or stainless-steel ferrules (2mm diameter, Precision Fiber Products, Chula Vista, CA) containing optical fibers (230um core, Thor Labs, Newton, NJ) were implanted bilaterally into the VTA (10° angle; from bregma AP: -6.0mm, ML: +2.3mm; DV: -7.1mm from dura). Implants were affixed to the skull using bone screws and dental acrylic, and ketoprofen (5mg/kg, sc) was administered during surgery and for up to two days thereafter, as needed, to reduce pain and swelling. At least 5 days were given for recovery prior to behavioral training, and at least three weeks were allocated for maximal expression of viral vectors.

### Chemogenetic control of neural activity

Inhibitory *designer receptors exclusively activated by designer drugs* (DREADDs) (AAV2-hSyn-HA-hM4D(G_i_)-IRES-mCitrine; Addgene, Watertown, MA) or control vector (AAV2-hSyn-EYFP; Addgene, Watertown, MA) were surgically injected bilaterally into the RMTg (400nL/side). The synthetic DREADDs ligand clozapine N-oxide (CNO, 1mM/0.5µl/side in 0.5% DMSO) or vehicle (0.5µl/side 0.5% DMSO) was injected 10mins prior to testing through chronic indwelling cannulae terminating bilaterally above RMTg axon terminals in the VTA. Intracranial injections were made by hand using a Hamilton syringe attached to water-filled PE50 tubing terminating in a stainless-steel injector (33 gauge, Plastics One, Roanoke, VA) extending 1mm past the implanted guide cannula. Injectors were left in place for at least 45sec to allow for drug diffusion. Each rat received two injections each of CNO or vehicle on separate days in a within-subjects counterbalanced design, and rats were given at least two days between injections to allow for drug washout.

### Punished cocaine seeking

We adapted our previously reported punished food-seeking paradigm [8] to a task where rats trained to self-administer cocaine, as described above, received once-daily sessions in which lever pressing for cocaine was immediately followed by presentation of light and tone cues, and brief (500ms) mild footshock. Each session began with an initial 5 “no cost” trials that were essentially identical to earlier training sessions, permitting individuals to “load up” and establish a steady rate of cocaine self-administration at the start of each test session. Beginning on the 6^th^ trial of every punishment session, responses on the active lever yielded cocaine infusion and brief footshock that gradually increased in intensity within session. In this way, shock intensity was relatively low in early trials (beginning at .25mA) but increased by ∼25% every 4 trials thereafter until rats effectively withheld responding for cocaine for at least 30mins or 2hrs had elapsed, whichever occurred first (for a timeline see Figure 1B). This procedure allowed for rapid and repeatable assessments of within-session shock tolerances that were generally stable from day to day. The maximum shock intensity subjects endured to receive cocaine is reported as the “shock breakpoint”.

**Figure 1.**
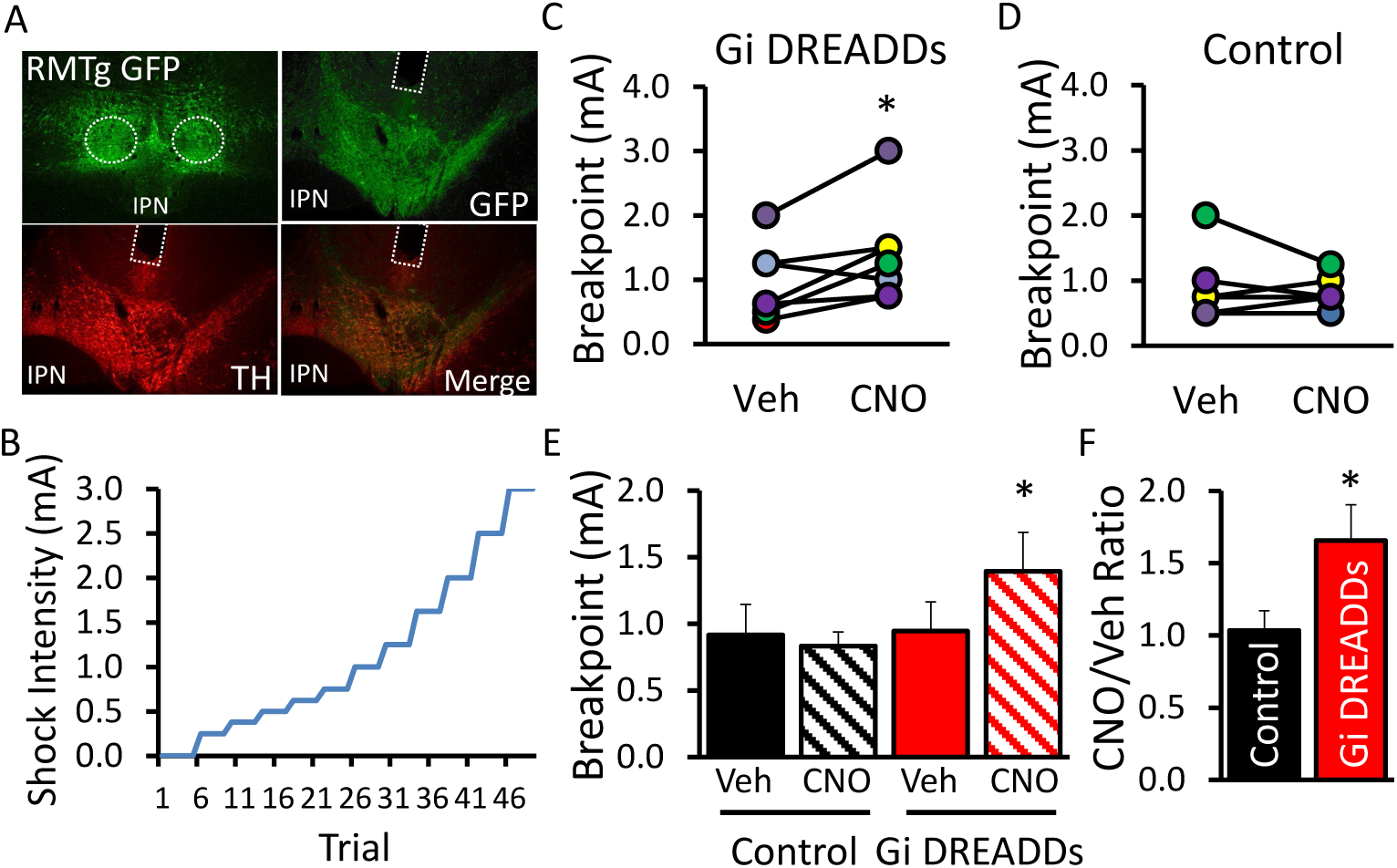
Chemogenetic inhibition of the RMTg→VTA pathway causes punishment resistance in lever pressing for cocaine. A) Rats received bilateral injections of inhibitory Gi DREADDs into the RMTg (upper left panel) and intracranial cannulae were implanted bilaterally above RMTg terminals in the VTA. B) Schematic of the punishment task where lever pressing for cocaine was punished by footshock that increased in intensity as the session progressed. (C-D) Individual data and (E) group means in shock breakpoint from rats expressing control vector or Gi DREADDs after pretreatment with either CNO (1mM) or vehicle (0.5% DMSO). F) Rats expressing Gi DREADDs showed a 65% increase in shock breakpoint after CNO versus vehicle sessions. Panel C: t(6)=-2.626, *p=0.039; Panel D: t(5)=0.542, p=0.611; Panel E: Drug (F1,11=2.437, p=0.147), Virus (F1,11=0.936, p=0.354); Drug x Virus (F1,11=5.188, p=0.044); n=6-7 per group; *p<0.05;

### Optical control of neural activity

Channelrhodopsin-2 (AAV2-hSyn-hChR2(H134R)-mCherry; UNC Vector Core, Chapel Hill, NC) or control vector (AAV2-hsyn-mCherry; Addgene, Watertown, MA), was injected bilaterally (400nL/side) into the RMTg and optical fibers were implanted bilaterally above RMTg axon terminals in the VTA. Blue light (447nM) was delivered via computer-controlled laser (Dragon Lasers, Changchun, China) mated to optical splitters (Precision Fiber Products, Chula Vista, CA) by an optical rotary joint (Doric Lenses, Franquet, Quebec, Canada). Blue laser light (20mW/side) was pulsed at a frequency of 50Hz (5ms duration).

### Operant food seeking

Rats expressing ChR2 or mCherry virus in RMTg with optical fibers in the VTA, or no surgery controls, were food restricted to ∼85% (±3%) of their free-fed bodyweight and trained to lever press for standard food pellets (45mg; LabDiet, St. Luis, MO) in daily sessions consisting of 70 trials or 60mins, whichever came first. Each trial began with the extension of both an active and inactive lever located on either side of a central food hopper, and a cue light was illuminated above each lever. Initial training was performed on an FR1 schedule that automatically incremented by one after every three consecutive trials completed in <15sec per trial up to a maximum FR5. Completion of the FR resulted in delivery of a single food pellet, both levers retracted, and the cue lights were extinguished, and a new trial commenced immediately thereafter. Once rats successfully escalated up to an FR5 over two consecutive sessions, the second phase of training commenced where each session was performed at a set FR5, while the other session parameters were identical to initial training. When rats reliably completed the FR5 sessions with low latency (>95% of trials completed in <15sec) for two consecutive sessions, subjects progressed to the third and final phase of training which was performed at a set FR5 with a 15sec ITI. After two consecutive sessions in this final training phase where >95% of trials were completed with <15sec latency, subjects were tested in once-daily sessions in which completion of the FR5 triggered the delivery of food and either brief mild footshock (0.5mA, 500ms duration) or pulsed blue laser light delivered bilaterally to RMTg terminals in the VTA. Rats expressing ChR2 received either 500ms or 30sec duration of optical stimulation with an additional 15sec ITI before the next trial to permit food consumption. Sessions terminated after subjects completed all 70 trials or after 60 min, whichever came first.

### Optical stimulation in cocaine seeking

Rats expressing ChR2 or control vector in the RMTg were implanted with optical fibers in the VTA, and an intrajugular catheter was implanted. Rats then were trained to self-administer cocaine as described above, at which point 4 additional self-administration sessions were given in which each lever press for cocaine was followed by optical stimulation of the RMTg→VTA pathway. The light-paired sessions were identical to self-administration training except a bilateral optical splitter was attached to the head, and a 2min timeout was imposed after completion of the FR during which time pulsed blue light was delivered to RMTg terminals in the VTA (See Figure 3A for a schematic of a representative trial). During the 2min of optical stimulation, the active and inactive levers remained extended and presses were recorded, but additional drug infusions or light/tone cues were unavailable. The duration of optical stimulation (2min) was longer than the light duration parameters used in the previous food-seeking paradigm; this was done to overlap and extend slightly past the period of cocaine delivery to account for differences in the timing and pharmacokinetics of intravenous cocaine infusion, which causes elevations in brain DA levels that persist for several minutes after IV infusion [20, 21].

### Histology

Subjects were deeply anesthetized by inhaled isoflurane and transcardially perfused using 0.9% saline followed by 10% formalin. Brains were removed and stored in 10% formalin overnight before being transferred to 20% sucrose with 0.05% sodium azide for cryoprotection. Tissue was collected in 40µm-thick sections on a cryostat or freezing microtome, and floating sections were processed using immunohistochemistry to verify virus expression in the RMTg and accurate placement of cannulae or optical fibers in the VTA. Tissue was incubated overnight in PBS with 0.25% Triton X-100 (Sigma-Aldrich) and primary antibody for rabbit anti-GFP (1:50K, Abcam), mouse anti-tyrosine hydroxylase (TH; 1:10K, Millipore Inc.), or rabbit anti-FoxP1 (1:20K, Abcam). Fluorescence was visualized using 30-min secondary incubation in either Alexa Fluor 488-conjugated donkey anti-rabbit or Cy3-conjugated donkey anti-mouse secondary (1:1000, Jackson Immunoresearch). Following each incubation step, tissue was rinsed 3X in PBS at 1min/wash. Data were included from animals showing GFP+ cells clustered bilaterally in the region corresponding to the FoxP1-positive RMTg [15], with dense labelling of axon terminals in the TH-positive VTA.

### Statistics

Student’s t-test was used to assess differences in shock breakpoint sorted by virus condition, and a repeated measures ANOVA was used to analyze mean shock breakpoints across groups. Ratios in shock breakpoints (mean CNO response divided by mean vehicle response) were determined for each individual animal and averaged across animals within each condition; a Student’s t-test was then performed on the mean shock ratios for each group. Repeated measures and between subjects ANOVAs were used to assess differences in lever pressing for food and latency to complete footshock or light-paired trials, as well as mean lever presses and infusions in cocaine self-administration and reinstatement studies. A two-way repeated measures ANOVA was used to assess differences in active versus inactive lever presses during self-administration training and light-paired sessions. Significant main or interaction effects were further probed using one-way ANOVA or student’s t-tests. When applicable, a Bonferroni correction was applied to adjust for multiple comparisons.

## Results

### RMTg→VTA activity is required to suppress cocaine use under punishment

Individuals battling addiction often discount the negative consequences of their drug use, choosing instead to continually engage in often dangerous and damaging patterns of behavior. Given our earlier findings suggesting a critical role for the RMTg→VTA pathway in suppression of seeking natural (food) reward under punishment, it was important to test whether this circuit is similarly involved in drug seeking when an aversive cost is incurred. Accordingly, rats expressing inhibitory (G_i_) DREADDs in the RMTg (Figure 1A) were trained to self-administer cocaine prior to testing in a novel punished drug-seeking task where lever pressing for cocaine is immediately followed by brief mild footshock that gradually increases in intensity as subsequent lever pressing for drug infusions occurs within session (see Methods-Punished Cocaine Seeking). A schematic of the paradigm is shown in Figure 1B. Pathway-specific inhibition of the RMTg→VTA circuit was accomplished through microinfusion of the DREADDs ligand CNO through cannulae aimed at RMTg terminals in the VTA 10min prior to test sessions.

In rats expressing inhibitory (G_i_) DREADDs, chemogenetic RMTg→VTA inhibition significantly increased the maximum shock rats endured to receive cocaine (p=0.039; Figure 1C). This effect was not otherwise explained by non-specific effects of CNO, as YFP-expressing controls did not exhibit changes in shock breakpoint between CNO vs vehicle treatments (p=0.611; Figure 1D). In contrast, inhibition of the RMTg→VTA circuit in rats expressing G_i_ DREADDs caused an ∼65% increase in the intensity of shock required to suppress cocaine use (Figure 1E-F). Notably, we have previously demonstrated that even complete ablation of the RMTg does not affect the sensory perception of shock, nor does it produce generalized learning impairments [8]. The present data, therefore, suggest that inactivation of the RMTg→VTA pathway leads to a compulsive pattern of persistent drug taking despite impending punishment.

### Optogenetic RMTg→VTA stimulation suppresses lever pressing for food reward

Given the above finding suggesting inhibition of the RMTg→VTA circuit causes persistent cocaine use under punishment, we next tested whether contingent stimulation of this pathway, in the absence of an overtly aversive stimulus, is sufficient to suppress seeking of natural (food) reward. To this end, rats underwent surgery for injection of virus encoding the excitatory opsin ChR2 and optical fibers were chronically implanted above RMTg terminals in the VTA (Figure 2A). Rats then were trained to lever press for food on an FR5 schedule of reinforcement prior to undergoing light stimulation sessions in once-daily sessions in which lever pressing for food was immediately followed by optical stimulation of the RMTg→VTA pathway (see Methods-Operant Food seeking). While contingent footshock (0.5mA, 500ms duration) in surgically naïve rats caused a significant reduction in total completed trials in each shock session (p<0.001; Figure 2B) and increased trial latency (p=0.03; Figure 2C), we found no change in food seeking when responding was immediately followed by optical RMTg→VTA stimulation of a comparable duration (500ms) (Figure 2B-2C).

**Figure 2.**
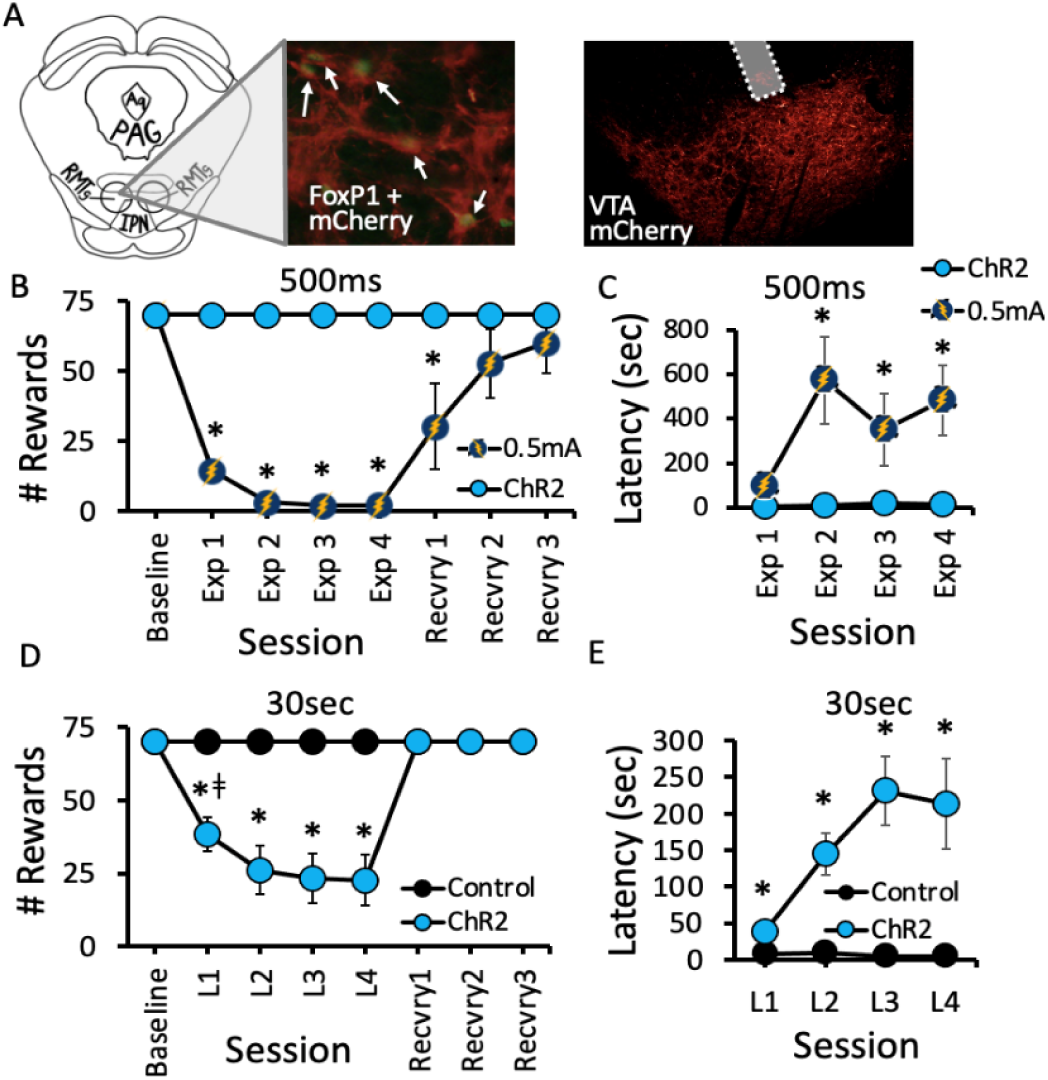
Optical RMTg◊VTA stimulation suppressed lever pressing for food reward**. A)** Rats received bilateral injection of virus encoding channelrhodopsin-2 (ChR2) or control vector in the RMTg (left), and optical fibers were implanted above axon terminals in the VTA (right). **B)** Contingent foot shock exposure (Exp) sessions 1-4; 0.5mA, 500ms duration) delivered immediately after successful completion of the FR significantly reduced the total number of trials completed compared to their pre-shock baseline and to 500ms optical RMTg◊VTA stimulation. In contrast, 500ms optical stimulation did not affect responding for food. **C**) After repeated footshock sessions, rats exhibited increased latency to complete food seeking trials when analyzed over the first 10 trials of each exposure session. **D)** Longer duration optical stimulation (30sec duration), however, robustly reduced the total number of food seeking trials ChR2 rats completed compared to mCherry controls. **E)** Rats receiving 30sec light exposure demonstrated increased latency to complete food seeking trials when analyzed over the first 10 trials of each test session. (Panel B: Session (F7,77 = 14.092, p<0.001); Condition (F1,15 = 206.389, p<0.001); Session x Condition (F7,88=12.304, p<0.001). (Panel C: Session (F4,32=3.076, p=0.03), Condition (F1,8=11.397, p=0.01), Session x Condition (F4,32=2.913, p=0.037). (Panel D: Session (F7,70=18.219, p<0.001); Virus (F1,10=22.747, p<0.001); Session x Virus (F7,70=18.219, p<0.001). *ǂ p<0.05 vs all other groups, *p<0.05 vs Baseline and L1. Panel E: Session (F3,27= 3.370; p=0.033), Virus (F1,9= 13.290; p= 0.005), Session x Virus (F1,9= 3.590; p=0.026). n=5-7 per group; *p<0.05 vs control.

To test whether more robust optical stimulation would be more effective at mimicking shock-induced suppression of food seeking, we generated a separate group of rats expressing either ChR2 or mCherry virus in the RMTg with optical fibers implanted in the VTA, and replicated the above experiment with 30sec duration contingent light delivery after each completed food seeking trial. While both ChR2-and mCherry-expressing rats completed the maximum 70 trials per session with short latency under pre-light conditions, 30sec RMTg→VTA stimulation significantly reduced the total number of trials completed over the 1-hour test session in rats expressing ChR2 (p=0.003; Figure 2D). Specifically, within the first light-paired session we observed a 46% suppression in total trials completed by ChR2-expressing rats. The effect of ChR2 photostimulation on food seeking became more pronounced with repeated testing such that by the 4^th^ light-paired session food seeking was reduced by 68% compared to controls. Assessing average trial latency over the first 10 trials of each light-paired session revealed that RMTg→VTA stimulation increased press latency on the active lever, an effect that similarly increased in magnitude across testing (Figure 2E). Notably, rats routinely were observed to consume food pellets immediately after cessation of light stimulation, and pellets were always absent from food trays at the end of each session, suggesting the manipulation did not act as a punisher for the food reward, per se, but rather for the previous action (lever pressing) that ultimately led to food delivery. Upon termination of photostimulation during “recovery” sessions 1-3 (Figure 2D), ChR2 rats readily resumed food-seeking behavior and once again completed the maximum 70 trials/session within the first recovery session without light.

### Optical RMTg→VTA stimulation concurrent with ongoing drug use produces long-lasting suppression of cocaine seeking

Given the finding that RMTg→VTA stimulation is sufficient to reduce lever pressing for natural rewards, we next tested whether stimulation of this circuit is also sufficient to reduce drug taking. Therefore, rats expressing ChR2 or control vector in the RMTg were trained to self-administer cocaine, prior to receiving 4 additional self-administration sessions where lever pressing for cocaine was paired with 2min of optical RMTg→VTA stimulation (Figure 3A). While initial variability was observed between groups in the early acquisition of cocaine self-administration, with control rats showing elevated pressing in early training relative to ChR2-expressing rats, responding rapidly equalized and remained stable with ChR2 and control groups showing similarly robust preference for the active (cocaine-paired) versus inactive lever (Figure 3B). A marked shift in behavior was observed, however, during the last 4 self-administration sessions (Light 1-4) when optical stimulation of the RMTg→VTA circuit occurred. During the photostimulation period when levers were still available, but cocaine and cues were withheld, we observed a significant increase in lever pressing in the control group that was conspicuously absent in rats expressing ChR2 (p<0.01, ctrl active presses vs all other groups). Indeed, controls showed a near tripling of presses during light-paired sessions while responding in the ChR2 group remained relatively stable at a level no different than pre-light self-administration in either the ChR2 or control groups (p=1.0; Figure 3C). This effect on lever pressing during light delivery did not translate to substantial changes in actual cocaine taking, however, as control rats did not take more cocaine during Light 1-4 sessions compared to their S8-10 baseline. Moreover, ChR2-expressing rats displayed no difference in mean cocaine infusions across light-paired sessions compared to their pre-light baseline, and we observed only a modest, but statistically significant, reduction in infusions during light-paired sessions in ChR2 vs control rats (p=0.044; Figure 3D). It is therefore likely that the reduced lever pressing observed in ChR2 rats during light-paired sessions primarily arose from a transient change in locomotor activity which has been reported elsewhere after manipulations of the RMTg [8, 12]. When the light terminated and a new trial was initiated, however, ChR2-expressing rats appeared to resume pressing for cocaine to a level no different than their pre-light baseline.

**Figure 3.**
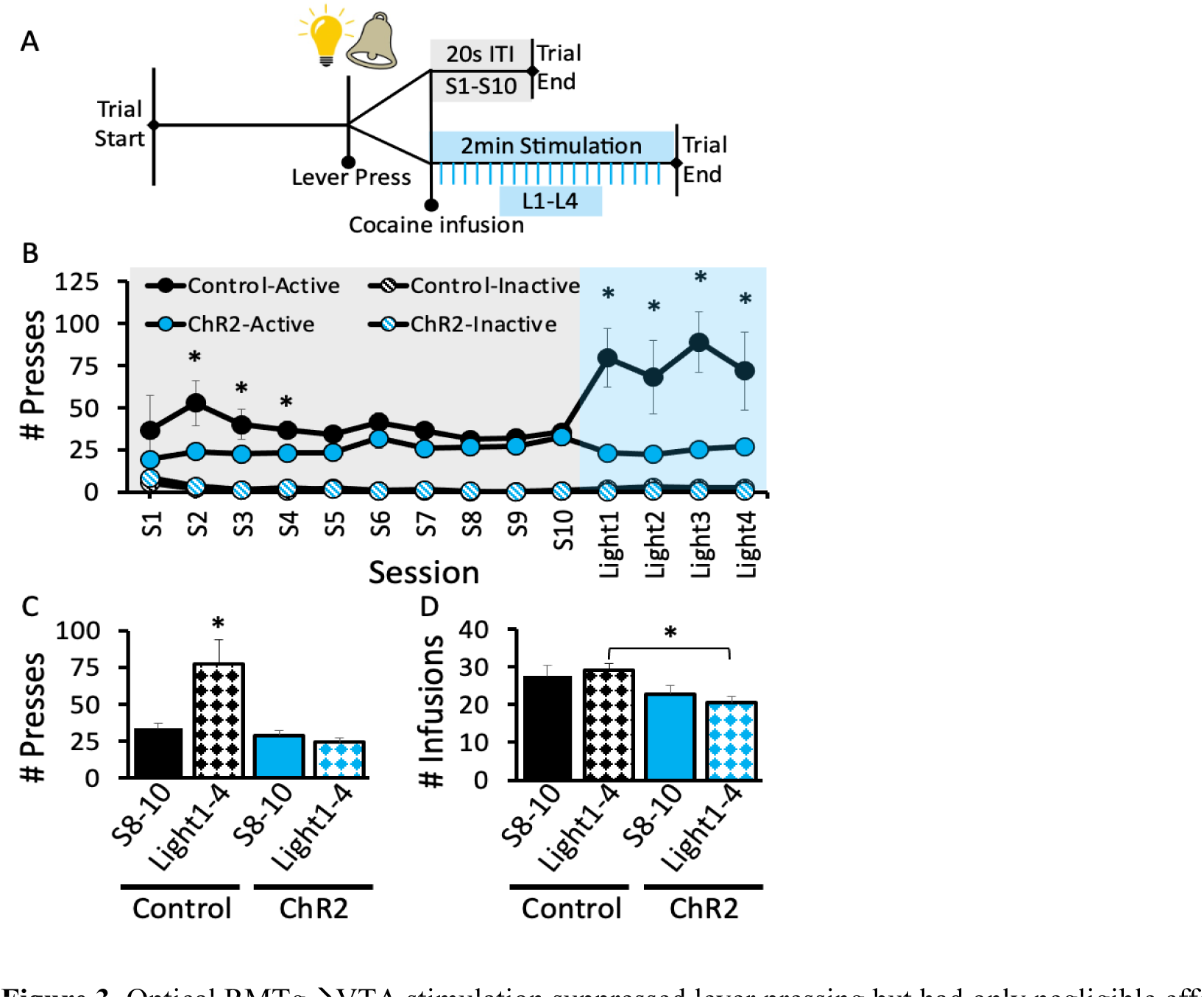
Optical RMTg◊VTA stimulation suppressed lever pressing but had only negligible effects on cocaine intake. Rats were trained to self-administer cocaine prior to receiving an additional 4 light-paired sessions in which lever pressing yielded light and tone cues, infusion of cocaine, and two minutes of optical stimulation (20mW/side, 50Hz with 5ms light duration). A) Schematic of trial in light-paired session. B) Rats in both the ChR2 and control groups rapidly acquired stable rates of cocaine self-administration that was nearly identical by the end of training. C) In light sessions 1-4, control rats displayed a significant increase in lever presses occurring during the extended intertrial interval in light-paired sessions. D) The number of earned cocaine infusions during light-paired sessions was relatively unaffected by optical stimulation, as both ChR2-expressing and control rats did not differ in the rate of infusions during light vs baseline (S8-10) sessions; however, a modest but statistically significant difference in infusions was found when comparing across groups during light-paired sessions. Panel B: Session (F2.86, 62.8= 4.239; p=0.01), Virus (F3,22= 33.209; p<0.001), Session x Virus (F8.56, 62.8 = 4.553; p<0.001). By S3 both ChR2 and control rats were significantly discriminating between active vs inactive lever (p<0.05). n=6-7 per group; *p<0.05 control active lever vs ChR2 active lever.

Taken together, the findings above indicate that while RMTg→VTA stimulation was sufficient to reduce lever pressing at the time of light delivery, this manipulation had only modest effects on disrupting cocaine use. Still, the possibility remained that some underlying change in the motivational value of cocaine had in fact occurred that we were unable to detect in FR1 responding for cocaine. We therefore extended testing after light session 4 (Figure 3B) to include a period of extinction training prior to testing for cue-induced reinstatement of cocaine seeking, a well-established model to test motivation to obtain drug [22]. Extinction behavior was remarkably similar in ChR2 and control rats; however, a profound difference in cue-induced reinstatement was observed (Figure 4B). While control rats exhibited robust reinstatement of lever pressing in response to the formerly cocaine-paired cues (p<0.001 versus Control and ChR2 Ext.10; Figure 4B), ChR2-expressing rats displayed a markedly blunted seeking response (p=0.012 versus Control Cue Test), emitting presses at a rate no different than extinction levels in the final session prior to the cue test (p=0.965 and p=0.904 versus ChR2 Ext. 10 and Control Ext.10, respectively). Also, despite the light-induced effects on lever pressing and number of infusions in ChR2 versus control rats (Figure 3C & D), those effects are likely independent of lever pressing during reinstatement, which did not correlate with either lever presses (Figure 4C) or drug infusions (4D) averaged across the 4 light-paired sessions. These data indicate that even though optical RMTg◊VTA stimulation hade only modest effects on reducing drug use when cocaine was readily available, the manipulation did result in long-lasting changes in the motivation to obtain cocaine when tested for cued reinstatement nearly two weeks later.

**Figure 4.**
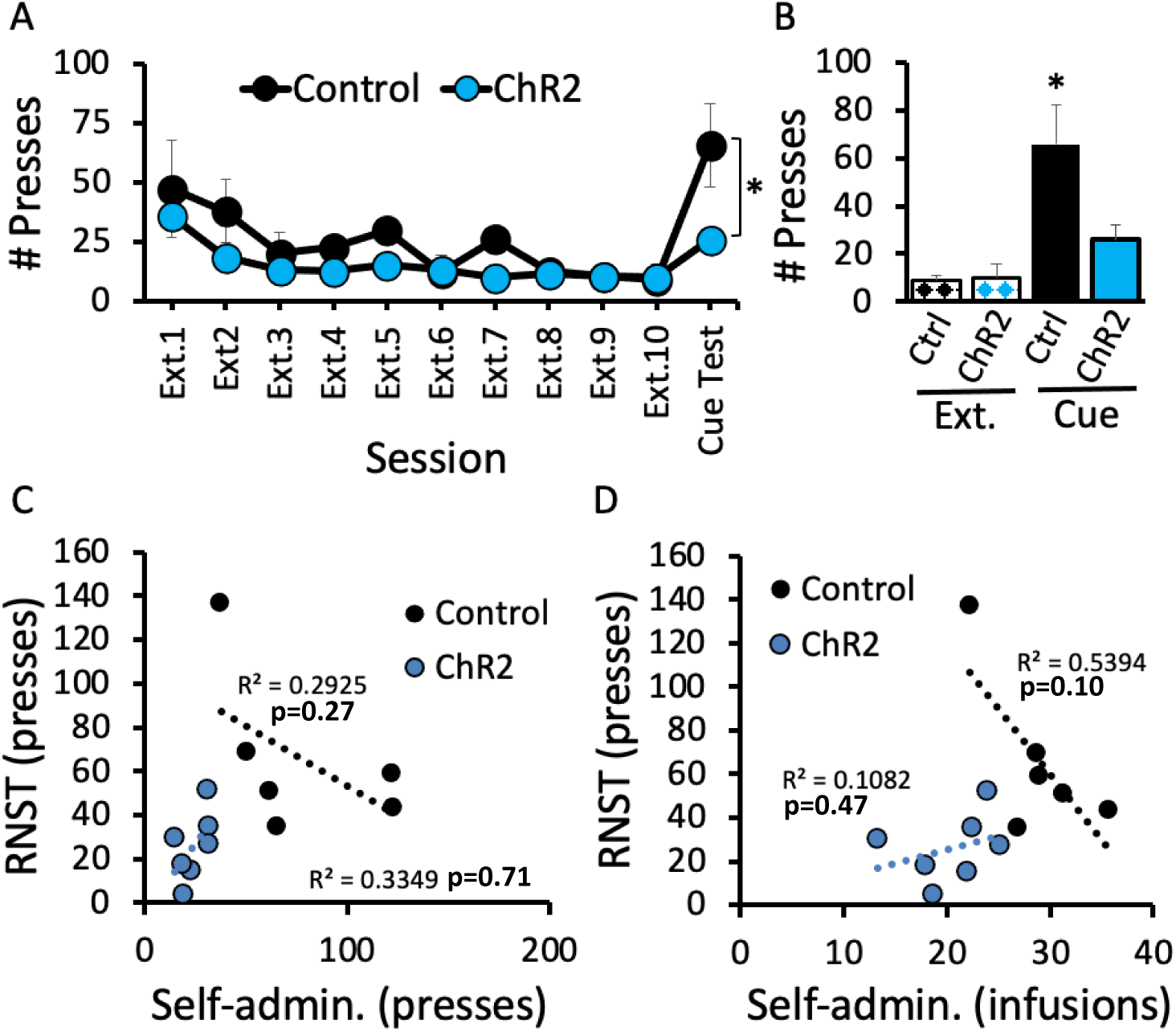
Cue-induced reinstatement was significantly attenuated by prior stimulation of the RMTg◊VTA pathway during cocaine self-administration, while extinction rates were similar between groups (A-B). Neither reinstatement lever presses nor infusions correlated with previous self-administration lever presses or infusions (C-D). Panel A: Session (F2.75,30.30= 9.624, p<0.001), Virus (F1,11= 6.89, p=0.024), Session x Virus (F2.75,30.30= 2.348, p=.097; Panel B: Group (F3,22=3.845, p=0.024), n=6-7 per group; *p<0.05 vs all other groups.

## Discussion

The ability to learn from and appropriately respond to the negative consequences of one’s actions is fundamental for survival [23], and deficits in aversive learning are observed across several neuropsychiatric disorders [24–26]. Here, we show that punishment-induced suppression of cocaine seeking depends upon signaling from the RMTg to the VTA, as chemogenetic inhibition of this pathway significantly increased the footshock intensity required to suppress responding for cocaine infusions. To recapitulate aspects of the neural “experience” to footshock in the absence of an overtly aversive stimulus, we optogenetically stimulated the RMTg→VTA circuit at the time when rats would have otherwise received footshock punishment, and found that this manipulation robustly suppressed lever pressing for food when delivered with 30sec, but not 500ms, duration. In contrast, optical RMTg→VTA stimulation during cocaine self-administration caused only a minor reduction in cocaine use when the drug was readily available, even at much longer duration stimulation that the food seeking experiments. Subjects nevertheless appeared to have learned something profound from this experience, however, as motivation to obtain cocaine was robustly suppressed when rats were later exposed to drug-associated cues nearly two week later, indicating enduring reductions in cocaine motivation.

Humans with a history of cocaine abuse display impairments in inhibitory control associated with alterations in prefrontal cortex activity [1, 3], perhaps underlying the decreased sensitivity to punishment observed in drug-addicted individuals [27–29]. Rodents also show addiction-like behavior after protracted drug use (∼3 months), with a subset (∼25-30%) developing insensitivity to the suppressive effect of footshock on cocaine self-administration [18, 19, 30–32]. Chen et al. (2013) found that this cocaine-induced punishment resistance is bidirectionally modulated by the prelimbic cortex, with optical stimulation restoring punishment sensitivity in shock-resistant mice while prelimbic inhibition caused punishment resistance in shock-sensitive individuals. While rats in the present studies were given only 10-14 days of short-access (2hrs) cocaine, which would otherwise be insufficient to cause punishment resistance [30, 31], RMTg inactivation nevertheless caused a punishment-resistant phenotype similar to that observed after protracted cocaine use. It is, therefore, important to note that the RMTg receives input from the prefrontal cortex (PFC) [33], and this pathway is required for both punishment-induced suppression of food seeking [9] and cue-induced reinstatement of cocaine seeking [34]; however, a role for prefrontal inputs to the RMTg in drug-induced punishment resistance has not been explored.

In contrast to the robust inhibitory effect of RMTg→VTA stimulation on food seeking, stimulating this pathway was largely ineffective at preventing ongoing drug use even despite the rather robust stimulation parameters used here. While *lever pressing* during light delivery in cocaine self-administration was indeed suppressed in ChR2 rats compared to controls, there was only a modest reduction in *cocaine infusions* between these groups, and when comparing within group effects on cocaine intake, optical stimulation did not cause a significant change from baseline in ChR2 rats. While the studies presented here are unable to discern the molecular or structural consequences of the different light parameters used in the food and drug experiments, it is possible that varying durations of photostimulation have different consequences on learning, motivation, and plasticity, irrespective of the specific reward or outcome. Alternatively, these differences may reflect different motivational values of natural versus drug rewards [33, 34], or the different molecular and pharmacological actions of cocaine versus food reinforcers on brain reward pathways [35]. Indeed, we would expect RMTg stimulation to inhibit DA cell bodies in the VTA given the relative selectivity of GABAergic RMTg efferents to DA neurons [36], while cocaine’s actions are primarily mediated by changes in DA neurotransmission [37]. It is unlikely, though, that the inability of optical stimulation to reduce drug taking arose from some non-specific limitation of our approach, as this manipulation did robustly suppress lever pressing during the ITI, an effect that is unsurprising given previously reported changes in locomotor activity after manipulations of the RMTg [8, 11]. What is surprising, however, is even though optical stimulation reduced lever pressing within the period overlapping light delivery, this effect was remarkably transient, as termination of the laser led to nearly immediate resumption of drug use as evidenced by the nearly identical number of infusions in pre-light self-administration versus light-paired sessions in ChR2-expressing rats. When abstinent from cocaine, however, drug-associated cues failed to induce vigorous responding, indicating some underlying change in the motivational value of cocaine or its predictive cues had occurred in response to the RMTg→VTA stimulation protocol. Notably, when lever pressing for food (Figure 2), normal responding rapidly reacquired after light-paired sessions concluded, indicating a somewhat nuanced interaction in learning from RMTg→VTA stimulation that may rely on several factors including reward availability and type of reinforcer (food vs drug).

Optical RMTg→VTA stimulation caused enduring effects that were revealed in several instances in the experiments described here. When lever pressing for food was followed by 30sec of RMTg→VTA stimulation, for example, rats completed fewer trials after repeated testing sessions, and the latency to complete trials at the start of each light-paired session became longer with repeated testing. Another learning effect, although more protracted, was on display when assessing cued reinstatement which was completely abolished when rats were tested weeks after pairing optical stimulation with cocaine self-administration. It is likely, therefore, plasticity induced by our stimulation protocol either disrupted communication of punishment-related signals from the RMTg to VTA, or rather altered communication from the VTA to downstream targets. It is notable that optogenetic stimulation of GABAergic afferents from the RMTg induces an adenylyl cyclase-dependent form of LTP and attenuates VTA DA firing [38]. Further, activity of GABAergic synapses onto the VTA is potentiated by several drugs of abuse, including morphine and ethanol [39, 40]. Notably, the various stimulation protocols used here did not seem to affect lever pressing in general, as we observed rats to reacquire lever pressing for food within a single session after light was terminated, and extinction behavior after the final light-paired self-administration session was indistinguishable between ChR2- and mCherry-expressing rats.

Animal models of maladaptive reward seeking have uncovered key insights into how brain circuits underlying features of addiction might be targeted to combat compulsive drug use [41]. Enduring treatments to prevent drug craving have been difficult to identify, however, and relapse rates remain high [2]. Here, we identify a critical circuit between the RMTg and VTA in suppressing reward seeking, [41] and demonstrate that RMTg→VTA stimulation serves as both an immediate punisher to reduce seeking natural reward, while also inducing long-lasting suppression of cued cocaine seeking during forced abstinence. Together, these data illuminate the RMTg→VTA pathway as a potentially fruitful target in producing enduring reductions in drug craving and relapse.

## Acknowledgements

The authors would like to thank the National Institutes of Health, NIDA Drug Supply Program for providing cocaine and CNO for these studies. We would also like to thank Grace Joyner for graciously providing artwork to the figures.

## Author contributions

Conceptualization and methodology: P.J.V. and T.C.J.; Investigation: P.J.V., D.P, S.L.B., and J.S.T; Writing: P.J.V. and J.R.W; Review and editing: T.C.J.; Funding: P.J.V. and T.C.J.

